# Fear Learning Induces Long-Lasting Changes in Gene Expression and Pathway Specific Presynaptic Growth

**DOI:** 10.1101/571331

**Authors:** Blythe C. Dillingham, Peter Cameron, Simon Pieraut, Leonardo M. Cardozo, Eun J. Yoo, Anton Maximov, Lisa Stowers, Mark Mayford

## Abstract

The stabilization or consolidation of long-term memories lasting more than a few hours requires new gene expression. While neural activity has been shown to induce expression of a variety of genes within several hours of learning, whether this leads to persistent changes in gene expression that lasts for days or weeks remains unclear. We developed a novel mouse line which expresses Cre recombinase in an inducible manner and used it to examine gene expression in learning-activated neurons of the medial prefrontal cortex (mPFC) one month following contextual fear conditioning. The mPFC is not required for the initial retrieval of contextual memory but becomes necessary after one month, suggesting a slowly developing plasticity. We found a variety of changes in gene expression in learning-activated neural ensembles that were specific to the mPFC. One group of transcriptional changes observed was the coordinated upregulation of presynaptic proteins suggesting a potential learning-induced elaboration of presynaptic terminals. We tested this idea by labeling the projections of mPFC neurons active during initial learning and found an increase in the number of terminals in neurons projecting to the basolateral amygdala at 1 month following training. These results suggest a presynaptic growth mechanism that could account for the enhanced role of the mPFC in fear memory retrieval at long time points after learning.

## Introduction

Memories undergo several phases of consolidation following learning. The stabilization of memory lasting 24 hours, a process known as synaptic consolidation, requires new gene expression^1,2^. Recently acquired episodic memories are dependent on the hippocampus (HPC) for retrieval but over time undergo a second process termed systems consolidation in which memory is believed to be incorporated into larger, distributed networks in cortical regions such as the medial prefrontal cortex (mPFC)^3–5^. In contextual fear conditioning (CFC) the neural ensembles encoding context in mPFC are recruited at the time of learning, but fail to support memory retrieval in the absence of the HPC at early time points^3,6–8^, suggesting a slowly developing form of plasticity in this cortical circuit. Current models postulate that systems consolidation involves the post-learning replay of hippocampal and neocortical ensembles representing the original learning experience that leads to strengthening of the neocortical representation allowing retrieval via the mPFC circuit at long post-learning timepoints (30 days in the case of CFC)^9,10^.

Although long-term memories have long been known to depend on new gene expression, the identification of specific functional genes mediating this process has been limited. High frequency neural activity induces the rapid and transient expression of immediate early genes (IEGs), such as the transcription factors cfos and zif268. These are necessary for late phase long-term potentiation (LTP)^11,12^ and the consolidation of context^11^ and spatial^13^ memories, however, the expression of these genes returns to baseline levels within several hours after neural activity. Two effector IEGs that directly impact plasticity and structural changes that accompany learning, Arc and Bdnf, also return to baseline quickly, within 24 hours after neural activity^14–19^. More recent studies of long-term memory have provided evidence for a role of epigenetic modifications such as histone acetylation^20–22^ and DNA methylation^23–25^, that may produce more prolonged alterations in gene expression.

To test whether learning produces prolonged changes in transcriptional profile, we developed a cfos-based genetic tagging system to examine the mPFC 30 days after CFC. We found that CFC produced a significant alteration in gene expression within learning-activated mPFC neurons at this timepoint. We found a variety of molecular pathways with altered expression, including a set of genes involved in synaptic vesicle release, which might reflect a learning-induced increase in presynaptic contacts. We tested this idea by introducing a synaptophysin-venus fusion protein into mPFC learning-activated neurons and found an increase in the number of presynaptic terminals in the basolateral amygdala (BLA) at 30 days relative to 7 days after training. This result suggests that at the time of learning, a long-lasting transcriptional program is activated, ultimately leading to an increase in mPFC-BLA connectivity that could account for the increased role of the mPFC in fear memory retrieval at remote timepoints.

## Results

In order to achieve neural activity-dependent genetic regulation we developed a novel mouse line in which a destabilizing domain Cre recombinase fusion protein (DD-Cre) is expressed under the control of the cfos promoter (FDC mouse). DD-Cre is comprised of a mutant *E. coli* dihydrofolate reductase (ecDHFR) fused to the N-terminus of Cre recombinase^26,27^. In the absence of the ecDHFR inhibitor trimethoprim (TMP), DD-Cre is unstable and degraded via the proteasome. In the presence of TMP, DD-Cre is stabilized and catalytically active (Figure 1A). In an effort to faithfully recapitulate endogenous cfos expression, the DD-Cre allele was inserted directly at the cfos translational start site, capturing both proximal and distal transcriptional regulatory elements^28^. We chose cfos as our locus of insertion as it is the most widely studied activity-dependent IEG that has been used to map neural activation in many behavioral paradigms and has been shown to label neurons that comprise a functional memory representation in CFC, sometimes referred to as engram neurons^29–32^. We used the recently developed DD-Cre for two reasons. First, TMP has been shown to rapidly cross the blood-brain-barrier and robustly activate DD-Cre after a single intraperitoneal (IP) injection^26^. Second, activity-dependent gene expression studies in memory have previously been limited to short time points or the use of drugs that may take weeks to clear the brain^33^ and activate endogenous receptors, potentially confounding behavioral results^34–36^.

**Figure 1.**
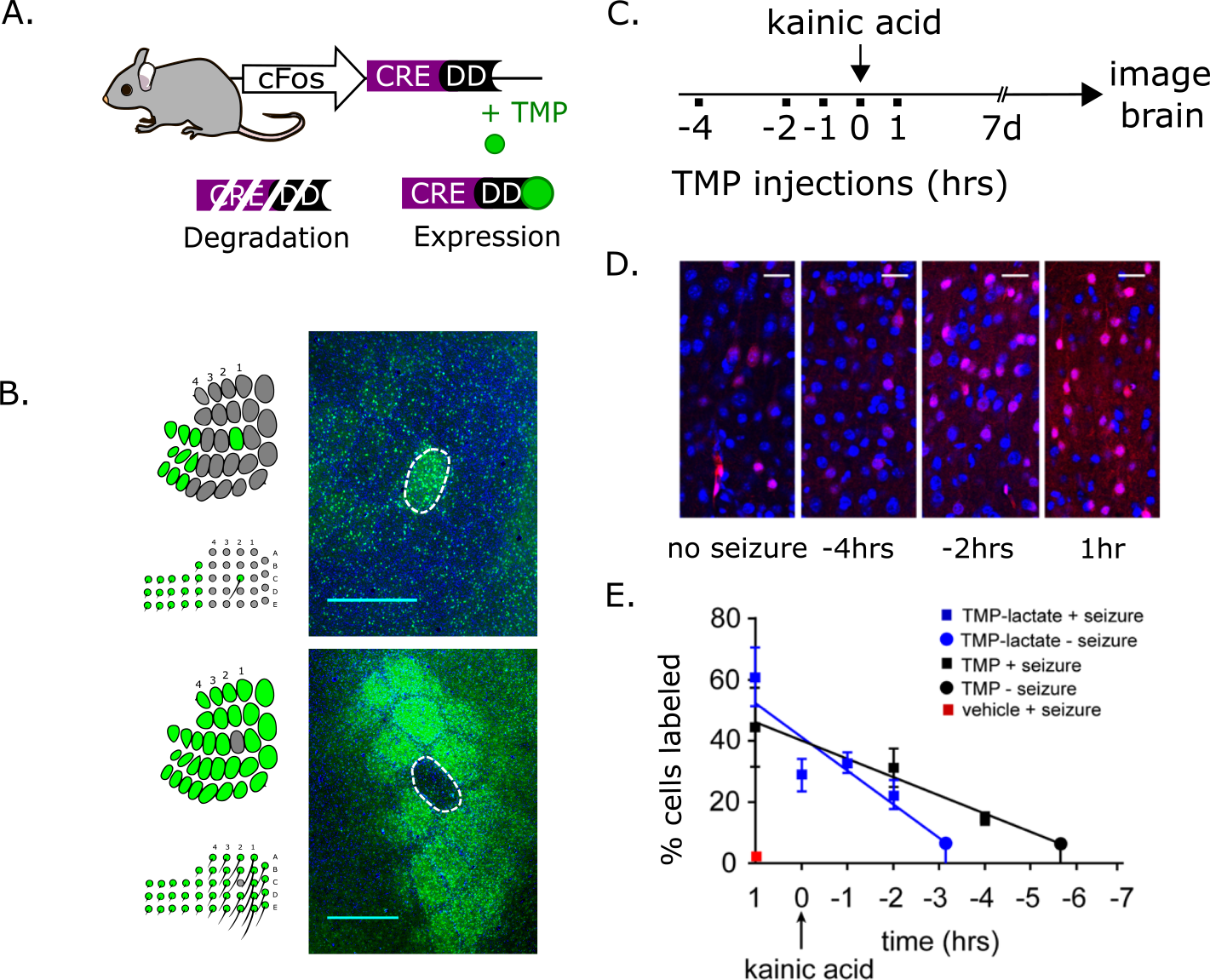
Destabilized Cre enables inducible, activity-dependent labeling system in mouse. A) FDC mouse schematic. DD-Cre was knocked directly into the cfos locus, replacing the translational start site. DD-Cre is stabilized and activated by the antibiotic trimethoprim (TMP). In the absence of TMP, DD-Cre is degraded via proteasomal pathway. B) Schematic (left) shows experimental design. Whiskers were either plucked surrounding the c2 position (top, left) or from the c2 position only (bottom, left). Representative image of c2 shows increased tdTomato labeling in the c2 barrel relative to surrounding barrels (top, right) (Green=tdTomato; Blue=DAPI), N=3. Representative image (bottom, right) showing decreased tdTomato labeling in the c2 barrel relative to surrounding barrels, N=3. Scale bar = 1mm. C) Experimental design to assess TMP/TMP-lactate clearance from the brain: TMP or TMP-lactate was injected at various time points relative to kainic acid. D) Representative images of layer 1 to layer 2/3 primary motor area from mice injected with TMP at various time points relative to kainic acid, or just TMP alone (no seizure) (Red=tdTomato; Blue=DAPI). Scale bar = 25μm. E) TMP and TMP-lactate clearance from the brain. Y-axis shows % cells labeled in primary motor area. X-axis shows time between kainic acid injection (t=0) and TMP/TMP-lactate injections. Mean ± SEM for each point seizure data point is shown. TMP best line fit (solid black line) intersects homecage “baseline” (black circle, 6.658 (mean) % cells labeled)” at 6.658 hrs, slope (−5.96 ± 2.05 [SE] % cells per hour) is significantly non-zero, p=0.0099 (F-test), r^2^=0.33; TMP-lactate best line fit (blue solid line) intersects homecage baseline (blue circle, 6.673 (mean) % cells labeled) at 4.142 hrs, slope (11.03 ± 3.00 [SE] % cells per hour) is significantly non-zero, p=0.0008 (F-test), r^2^=0.29. Leak due to seizure is minimal, vehicle + seizure mean ± SEM = 2.33 ± 0.563 % cells labeled.

To characterize the properties of the FDC line, we generated mice that were heterozygotes for both FDC and the Ai9 Cre reporter line (FDC-Ai9). Cells that have undergone Cre recombination express tdTomato, with native fluorescence readily detectable 2-3 days after the recombination event^37^. We investigated background tdTomato expression in FDC-Ai9 mice in the absence of TMP in 24 brain regions and found sparse labeling with about 1-3% of total cells labeled at 4 weeks of age (Supplemental Table 1). An exception was the granule cells of the dentate gyrus and the main olfactory bulb, which showed significant leak, suggesting that these cells express high levels of cfos at some point throughout development, or alternatively that DD-Cre is not processed and degraded equivalently in all cell types.

To determine the extent to which FDC mice report neural activity, we asked whether we could label cells in the primary somatosensory cortex (SSp) by whisker manipulation. The mouse SSp is organized into “whisker barrels,” functional units that receive afferent information mostly from a single contralateral vibrissae^38^. Previous experiments have shown that movement of individual whiskers induces cfos expression in the cognate contralateral whisker barrel^39–41^. We tested this in FDC-Ai9 mice using two whisker removal treatments: we plucked whiskers surrounding the c2 position, or conversely, we plucked only the c2 whisker and left the surrounding whiskers intact (Figure 1B). For each mouse, we left the whiskers on the opposite side intact to serve as a control. After plucking, mice were moved to a new cage with an "enriched" environment, which included a variety of novel objects. One day later, mice were injected with TMP, and brains were subsequently processed two weeks later to assay for differential whisker barrel labeling. Tangential sections revealed that mice with the c2 whisker left intact showed a dramatic increase in c2 barrel labeling relative to the surrounding barrels (Figure 1B, Supplemental Figure 1A&B). Similarly, plucking only the c2 whisker produced a significant decrease in tdTomato labeling in the c2 barrel relative to surrounding barrels. The whisker barrels contralateral to the unperturbed side showed uniform labeling (Supplemental Figure 1A). Thus, the FDC driver mouse line can be used to induce recombination specifically in neural ensembles responsive to natural sensory stimuli.

In addition to cfos activity and kinetics, the length of time the stabilizing ligand is active in the brain will also influence the temporal resolution of activity-dependent FDC driven labeling. To assess the functional time course of TMP-induced recombination, we injected TMP at 75μg/g, its lowest effective dose (Supplemental Figure 1C,D) at multiple time points relative to a seizure inducing stimulus (Figure 1C,D) and found the highest tdTomato labeling if TMP was given 1 hour post-seizure induction with diminished labeling at later time points. Linear regression analysis suggested that active TMP cleared the brain 6.7 hours after injection (95% confidence interval of clearance after injection = 4.7 to 16.9 hrs) (Figure 1C-E). We tested an additional ecDHFR inhibitor, TMP-lactate, a soluble derivative of TMP, and found similar results but with a slightly faster clearance relative to TMP (95% confidence interval of clearance after injection = 4.1 to 7.6 hrs) (Figure 1E). Thus, the time window for intraperitoneally injected TMP and TMP-lactate induced Cre activation in the FDC mice appears to be on the order of several hours. Due to its solubility and faster clearance we used TMP lactate for all subsequent experiments.

To test for transcriptional alterations in neurons activated during CFC, we crossed the FDC mouse to the RiboTag mouse^42^ (FDC-Ribo mice), which expresses an HA-tagged ribosomal protein RPL22 in a Cre-dependent manner, to allow inducible ribosome tagging in cfos active neural ensembles for subsequent immunoprecipitation and isolation of active mRNA transcripts as described previously^42^ (Figure 2A). To verify TMP-induced HA expression in FDC-Ribo mice, we fear conditioned animals on day 1 and 2 and injected TMP (FC+TMP) or saline (SAL) 10 minutes after each training session. Mice were fear conditioned two days in a row to maximize HA induction and provide an accurate measure of learning through freezing levels tested during the 3 minute pre-shock period on day 2. Thirty days later, we observed virtually no HA positive cells in the mPFC in SAL animals relative to FC+TMP, indicating tight regulation of our system (Figure 2B,C). This result was not due to differences in learning as both FC+SAL and FC+TMP froze at equal levels 24 hours after learning (Figure 2D). We also injected homecage control animals with TMP (HC+TMP) and observed HA expression levels significantly greater than SAL animals but less than the FC+TMP group (Figure 2B). Previous studies using cfos-based tagging systems have demonstrated that neurons active during learning in both cortical and subcortical regions are subsequently reactivated during retrieval and sufficient to drive behavior when optogenetically stimulated^7,30,31,43^. To confirm reactivation during retrieval of the learning-activated neural ensemble in the mPFC, we fear conditioned mice and injected TMP (Figure 2E) to label active neurons and performed a memory retrieval test 30 days later followed by immunostaining for HA (learning tag) and endogenous cfos (retrieval tag). Immunostaining indicated significantly more cfos expression in the 30 day retrieval group (RET) relative to a control group that did not receive the retrieval trial (HC), consistent with previous studies showing engagement of the mPFC in CFC memory retrieval at this time point (Figure 2H)^3^. In addition, we found significantly more reactivation in the RET group as evidenced by coexpression of HA and cfos (Figure 2F,G), consistent with recruitment of the original learning-activated ensemble during memory retrieval. Interestingly, this result was specific to mPFC as there was no significant difference in reactivation levels between RET and HC groups in the SSp (Figure 2G).

**Figure 2.**
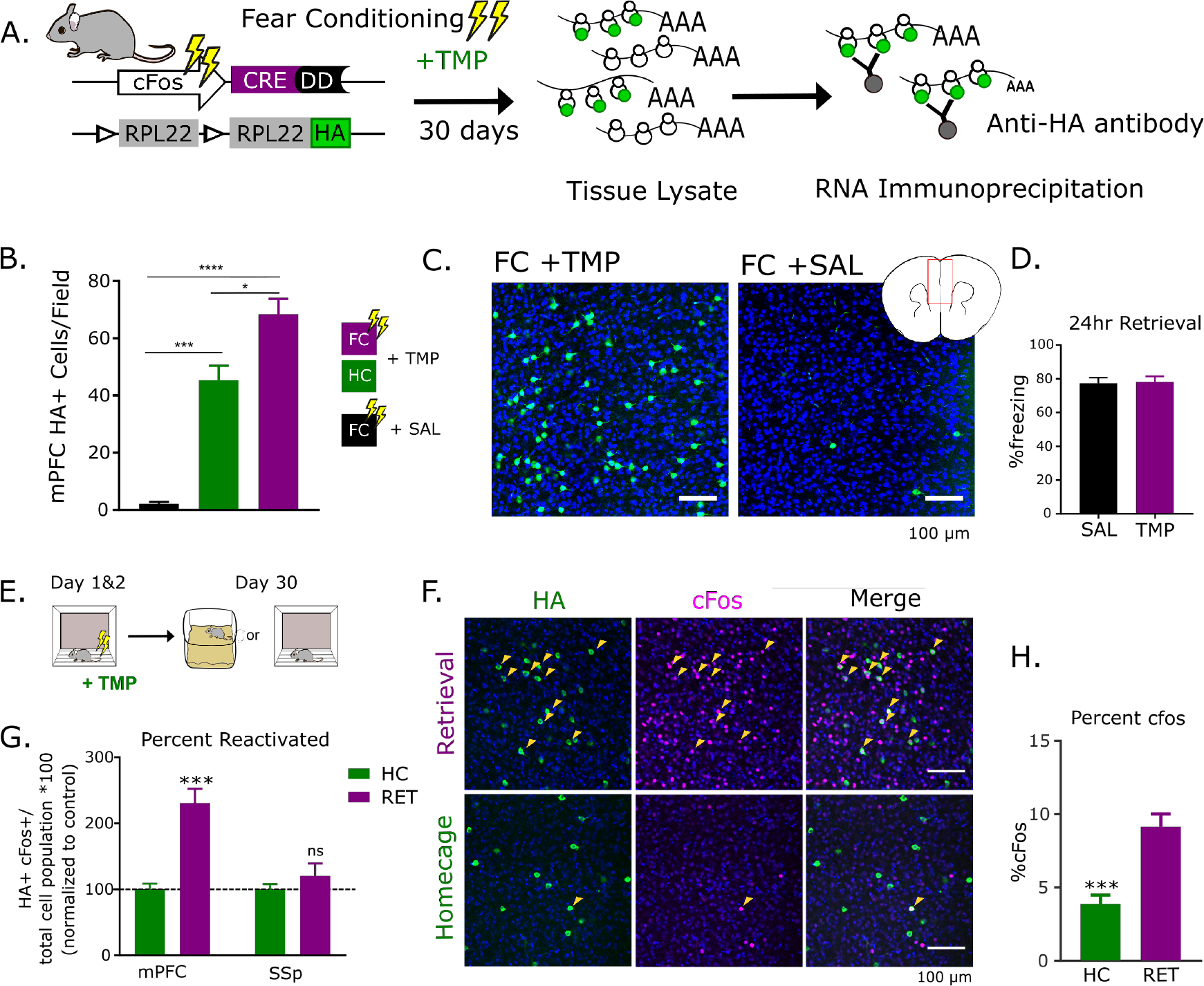
Learning-activated mPFC neurons are reactivated during 30 day memory retrieval in FDC-Ribo mice. A) FDC-Ribo mouse schematic. Upon TMP injection, HA-tagged ribosomal protein RPL22 is induced in active neurons allowing for subsequent RNA isolation from HA-tagged ribosomes. B) Significantly more mPFC HA cells/field in FDC-Ribo mice fear conditioned with TMP injection (FC+TMP) relative to saline injected controls (FC+SAL). Homecage controls injected with TMP (HC +TMP) showed significantly less mPFC HA cells/field relative to FC+TMP but significantly more than FC+SAL. N= 5-8; Mean ± SEM; Tukey’s multiple comparison test *p<0.05, **p<0.01, ****p<0.0001 C) Representative images HA expression in mPFC (Green=HA; Blue=DAPI). D) No significant differences in freezing levels during 24hr retrieval in FC+TMP or +SAL groups. E) Experimental design. Mice were injected with TMP and 30 days later given a retrieval trial or left in their homecage (HC). F). Representative images of HA and cfos staining in mPFC of HC and Retrieval (RET) groups. Green=HA; Magenta= cfos; Blue = DAPI.G) FDC-Ribo mice exhibited significantly more reactivation (HA co-expression with cfos) of mPFC ensembles during RET relative to HC. Reactivation normalized to total cell population [(HA+cFos+ cells)/(HA+ + cFos+ - HA+cFos+)] and shown as percentage of control. N=5-7; Mean ± SEM; unpaired t-test with Welch’s correction ***p<0.001. No significant differences were observed between groups in the somatosensory cortex (SSp). N=4. H) cfos expression was significantly higher in RET group relative to HC in mPFC. Mean ± SEM Unpaired t-test ***p<0.001.

To further characterize these active ensembles, we performed immunostaining in both FC+TMP and HC+TMP animals (Figure 3A) for the neuronal marker, NeuN, and a marker for inhibitory neurons, Gad65/67. We found 94% co-expression of HA and NeuN in both the FC and HC groups (Figure 3B,C). Consistent with previous experiments using cfos-driven systems in fear conditioning^29,44^, these cells were largely excitatory as indicated by only 2.3% and 3.4% overlap with Gad65/67 in FC and HC groups, respectively (Figure 3B,D).

**Figure 3.**
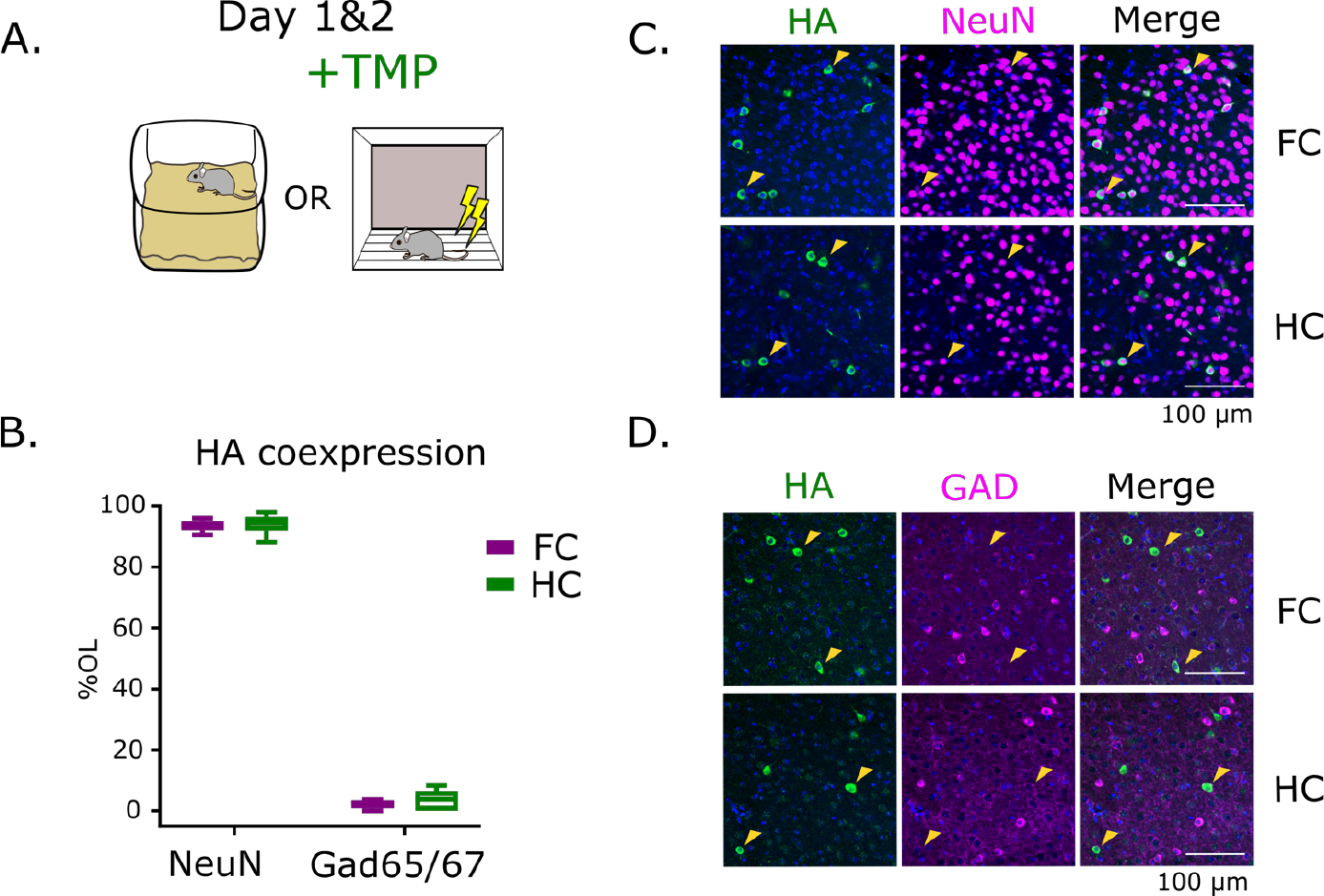
mPFC active ensembles are mostly excitatory neurons. A) Experimental design. FDC-Ribo mice were injected with TMP in their homecage (HC) or fear conditioning (FC). B) No significant differences observed between HC and FC groups in NeuN or Gad65/67 expression. HA coxpression with NeuN = 93.66% (FC) and 94.03% (HC). HA coexpression with Gad65/67 = 2.253% (FC) and 3.379% (HC). N=5-7. C) Representative images of NeuN (Magenta) overlap with HA (Green); Blue = DAPI D) Representative images of Gad6/67 (Magenta) overlap with HA (Green); Blue = DAPI.

To examine long-lasting transcriptional changes within learning-activated mPFC neurons, FDC-Ribo mice were injected with TMP after FC or HC, and 30 days later, we dissected the mPFC, purified HA-tagged ribosomes, isolated RNA and performed RNAseq analysis (Figure 4A). We found 1500 differentially expressed genes of which 700 were upregulated in the FC group (Figure 4B). To identify enriched pathways in FC mice, we used the DAVID open source analysis tool^45^ and found the most significant upregulated pathways were oxidative phosphorylation, neurological disease related, the proteasome, and the synaptic vesicle cycle (Supplemental Figure 2A). Taken together, this suggests that learning-activated mPFC neurons exhibit increased metabolic activity and presynaptic gene expression. As seen in Figure 4C, within our data set are genes that encode for proteins in each step of the synaptic vesicle cycle. Additionally, the changes in transcriptional profile are associated with multiple neurological disorders that may implicate memory or memory related processes^46^, and the upregulation of the proteasome pathway is relevant as recent studies have demonstrated a role for protein degradation in the maintenance of long-term potentiation (LTP) and in context fear memory retrieval^47,48^.

**Figure 4.**
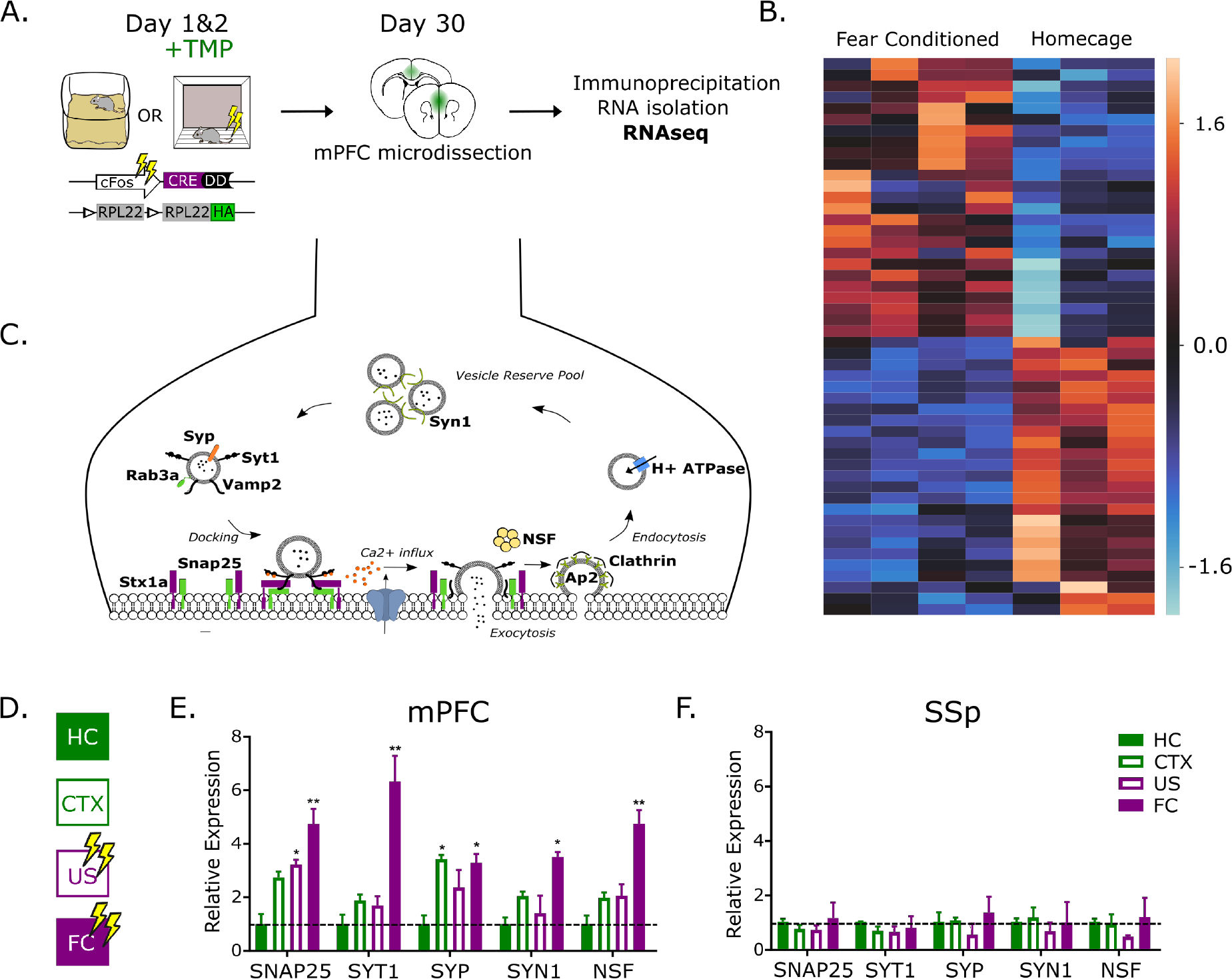
Contextual fear conditioning induces long-lasting increases in presynaptic gene expression in learning-activated mPFC neurons. A) Experimental design. FDC-Ribo animals were injected with TMP during fear conditioning (FC) or in their homecage (HC) and the mPFC was removed 30 days later after which immunoprecipitation of HA-tagged ribosomes and subsequent RNA isolation and RNAseq were performed. B) Heatmap shows z-scores (of log2 counts per million) of a representative sample of the differentially expressed genes (one for every 30); N=3-4; Each row = gene; each column = mouse. C) Schematic showing synaptic vesicle cycle genes (in bold) enriched in FC mice. D) For RNAseq validation, new biological replicates were used and two control groups were added. Immediate shock (US) and context exposure alone (CTX) in which minimal fear memory is acquired. E) qPCR validation of representative sample of presynaptic proteins enriched in FC mice. SNAP25 was significantly enriched in CTX, US and FC groups relative to HC control (*p=0.02 **p=0.002). SYT1 (*p=0.01), SYN1(*p=0.01) and NSF (**p=0.0018) were significantly only enriched in FC relative to HC. SYP was significantly enriched in both CTX (*p=0.049) and FC (*p=0.014) relative to HC. N=4; Mean ± SEM. Dunnett’s multiple comparison test. All asterisks represent significance relative to HC group. F) No significant differences found between FC, CTX and US relative to HC in the somatosensory cortex (SSp). N=4; Mean ± SEM.

To validate our RNAseq data, we repeated the experiment with new biological replicates and confirmed the upregulation of a representative sample of the synaptic vesicle cycle proteins by qPCR (Figure 4D,E). In addition, we asked if these gene expression changes are specific to associative learning by isolating RNA from mPFC neurons active during context exposure alone (CTX) or an immediate shock paradigm (US), both of which result in significantly less freezing than CFC (Supplemental Figure 2B,C). Those genes upregulated specifically in the CFC group included the calcium sensor synaptotagmin (Syt1), the vesicle membrane protein synapsin (Syn1), which binds to actin and sequesters vesicles in the reserve pool^49^, and the ATPase Nsf, which is involved in vesicle fusion at the presynaptic membrane^50^ (Figure 4E). Lastly, we confirmed mPFC specificity of these gene expression changes by performing qPCR on SSp active neurons under the same conditions (Figure 4D,F) and found no significant alterations in gene expression in the cfos labeled neurons from this brain region.

We next asked whether the enrichment of presynaptic proteins was indicative of an increase in synapse number and also sought to determine the time course of these changes. If there is synaptic growth, does this happen shortly after learning or slowly over time as memory is consolidated? To address this question, we microinjected an AAV to express a Cre-activated synaptophysin-venus fusion protein into the mPFC of FDC mice to label axon boutons in active neurons (Figure 5A,B). To normalize for variation in virus infectivity, mice were coinjected with a Cre-activated nuclear restricted fluorescent reporter, as it is difficult to reliably quantify cell bodies with the synaptophysin-venus label. Where distinguishable, however, the two viruses demonstrated complete overlap in Cre activated expression (Figure 5C). As outlined in Figure 5D, we used two groups of mice injected with TMP during CFC training and sacrificed 7- or 30-days later (7d post-FC and 30d post-FC), as well as two groups of homecage controls labeled in a similar manner (7d HC and 30d HC). In order to control for any background Cre recombination, the length of time from AAV injection to sacrifice was 38 days for all groups. We examined axon boutons in the BLA as it is known to have reciprocal connections with the mPFC and is implicated in the expression of learned fear^51^. The role of the BLA in remote memory remains largely unexplored, but lesion studies have shown its necessity for both auditory and context fear memory retrieval even 1.5 years after training^52,53^. We found approximately twice as many axon boutons/μm^2^ from mPFC projection neurons in the BLA 30 days after fear conditioning relative to the 7 day group (Figure 5E), suggesting that a slow progression of presynaptic growth occurs along the same timeline as memories are consolidated and rendered independent of the hippocampus. We found this result to be specific to the BLA projections, as we did not observe any significant differences in the number of mPFC to striatum terminals (Figure 5F). This result was not due to differences in learning as both fear conditioned groups froze at equivalent levels during the final retrieval trial (Supplemental Figure 3A), nor was it a result of synpatophysin-venus accumulation over time as the number of boutons in 30d HC and 7d HC were not significantly different from each other. Interestingly, we also analyzed synaptophysin-venus signal at a lower magnification by quantifying the area of signal/total area of the BLA (μm^2^) (Supplemental Figure 3B,C) and found that the lower magnification analysis correlated significantly (Supplemental Figure 3D) with the boutons/ μm^2^ data, providing an alternative form of analysis in instances where higher magnification is inaccessible.

**Figure 5.**
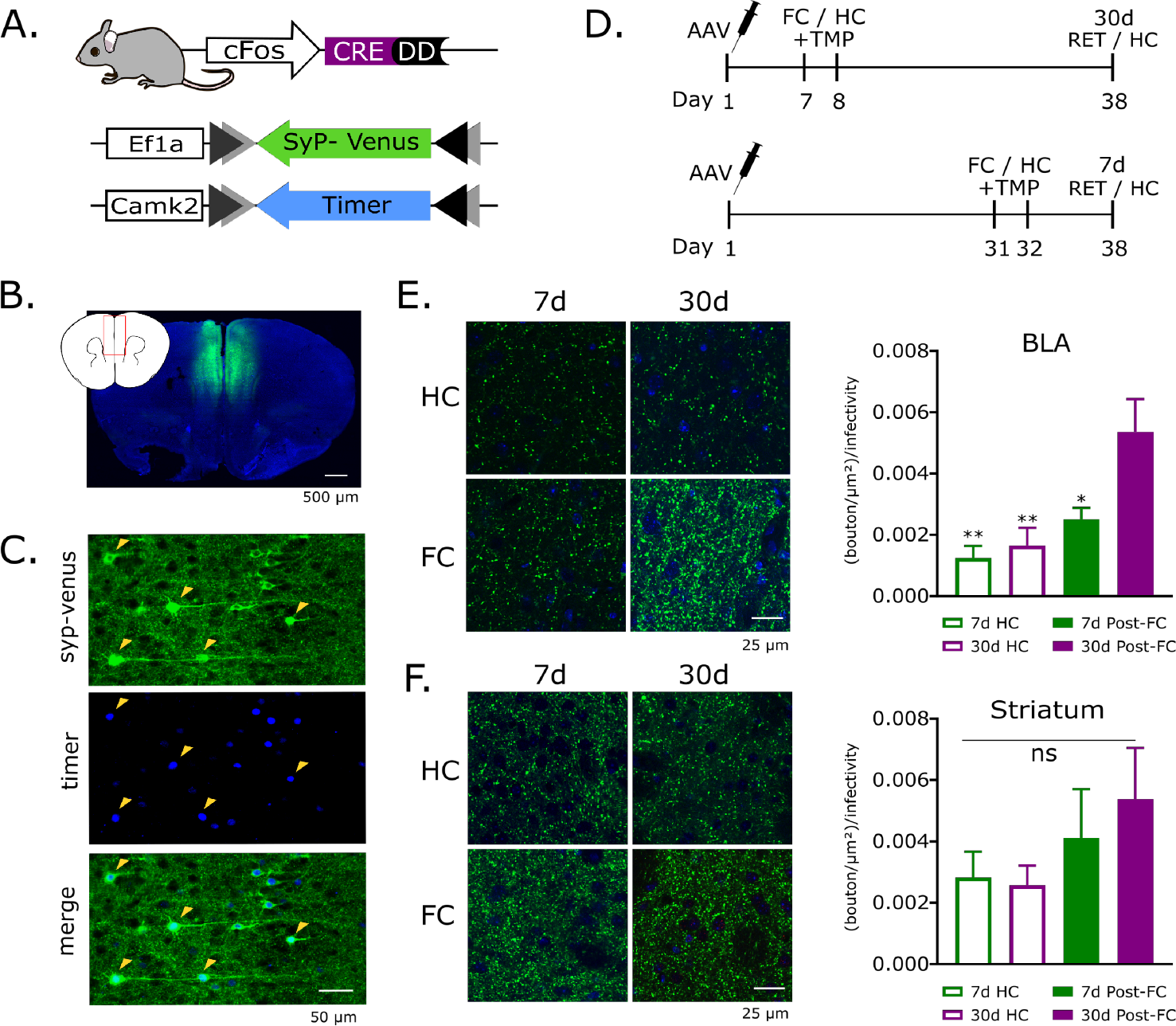
Increased presynaptic growth occurs in learning-activated mPFC neurons projecting to the basolateral amygdala 30 days after learning. A) FDC Mice were injected in the mPFC with an Ef1a-driven cre-activated synaptophysin-venus AAV in combination with a Camk2-driven cre-activated fluorescent timer reporter. B) Synaptophysin-venus expression (green) in mPFC; Blue=DAPI. C) Representative images showing synaptophysin-venus (green) overlap with timer (blue). D) Experimental design. E) Significantly more axon boutons/μm^2^ from learning-activated mPFC neurons in the basolateral amygdala (BLA) 30 days relative to 7 days after FC (*p=0.0329) as well as relative to HC controls at 7 (**p=0.002) or 30 days (**p=0.008). Boutons/μm^2^ for all groups are normalized to percentage of infected cells in mPFC. N=5-6; Mean ± SEM; Tukey’s multiple comparisons test. All p-values are relative to 30d post-FC group. Green=Syp-venus; Blue = DAPI F) No significant differences in boutons/μm^2^ in learning-activated mPFC neurons that project to the Striatum 30 days relative to 7 days after fear conditioning (p=0.8926) nor in HC controls at 7 day (p=0.5385) or 30 day (p=0.4234). Boutons/μm^2^ for all groups are normalized to percentage of infected cells in mPFC. N=5-6; Mean ± SEM; Tukey’s multiple comparisons. All p-values are relative to 30d post-FC group. Green=Syp-venus; Blue = DAPI.

## Discussion

The formation and consolidation of memories lasting days or longer requires new gene expression^2^. Although prevailing theory posits that memories are initially encoded in the HPC and become more dependent on the neocortex over time, recent studies support the role of cortical ensembles, including the mPFC, in the initial encoding of memory^7,54,55^. This raises the question, what are the molecular mechanisms underlying the transformation of hippocampal-dependent to hippocampal-independent memories within the mPFC? In the current study, we addressed this question using a newly developed transgenic mouse in combination with genome-wide RNAseq and found changes in transcription levels in learning-activated mPFC ensembles 30 days after training, with a specific increase in synaptic vesicle proteins. We then used genetic labeling of these activate mPFC ensembles with a synaptophysin-venus fusion protein to identify presynaptic terminals and found significantly more axon boutons in the BLA 30 days after fear conditioning relative to 7 days. Taken together, our data suggest a slow growth process in BLA-projecting mPFC neurons recruited at the time of learning and corresponding to long-lasting learning-induced changes in gene expression.

Memory formation and storage through learning-induced structural and functional synaptic changes has been postulated since the early descriptions of neuronal anatomy. Indeed, structural changes that occur after learning or LTP have been primarily linked to the formation or modification of dendritic spines in the HPC^56–58^. In cortical regions, one study used a similar cfos-based tagging system and demonstrated an increase in spines in active mPFC ensembles at 12 days relative to 2 days after CFC^7^. Also, consistent with systems consolidation, two studies have shown learning-induced spine growth in the anterior cingulate cortex (ACC) region of the mPFC increases over time, with the opposite effect seen in the HPC^59,60^

There are surprisingly few reports linking learning to presynaptic growth. In the mossy fiber pathway, which innervates the CA3 region of the HPC, synaptogenesis has been reported at both 24 hours and 7 days after training of a spatial memory task in rats^61,62^. Learning-induced presynaptic growth also been reported in the gill-withdrawal reflex of *Aplysia californica*. Electron microscopy studies demonstrated an increased number of active zones and presynaptic varicosities in sensitized animals immediately after learning that lasted for 3 weeks, paralleling the time course of memory retention^63–65^. This result was dependent on both protein and RNA synthesis^66^ and suggests that an increase in synapse number contributes to the persistence of memory. Another study looked at the mPFC and found that CFC induced a rapid reorganization of synaptic structure, including an increase in synaptic vesicles and larger active zones, just one hour after learning^6^. Our findings diverge from these studies in that we observed an increase in presynaptic terminals that develop over the course of 1 month. To our knowledge, just one study, in a social transmission of food paradigm^54^, found an increase in the presynaptic protein synaptophysin in the orbitofrontal cortex 30 days but not 1 day after learning. While this progression of presynaptic growth is consistent with our model of slowly developing plasticity into cortex, it lacks circuit specificity to memory relevant neurons.

CFC and other learning paradigms induce transient IEG expression which is believed to initiate a downstream transcriptional program that ultimately leads to permanent memory storage. Interestingly, blocking of Arc and Bdnf expression in the HPC 12 hours after learning^18,19^, outside of the standard time window to disrupt synaptic consolidation, impaired memory 7 but not 2 days later, suggesting a role for second a wave of gene expression involved in memory persistence. Still, these expression levels returned to baseline within 24 hours, indicating the persistence was likely a result of downstream molecular mechanisms. How then are memories maintained? Epigenetic modifications provide an appealing mechanism for instituting long-lasting changes in gene expression. In fact, one study found that inhibiting DNA methylation in cortex 30 days after CFC resulted in memory impairment 1 day later, suggesting that persistent DNA methylation helps maintain cortical memories long-term^24^. More studies are needed, however, to further understand this process.

Next generation sequencing technologies provide a unique opportunity to investigate genome-wide changes in gene expression in a hypothesis generating manner. Here, we have taken advantage of the cfos promoter and RNAseq to assess long-lasting changes in gene expression, specifically in mPFC learning-activated ensembles. Previous studies have performed transcriptional profiling in learning-activated neurons or whole brain regions after CFC but were limited to short-time points^6,67–69^. Our results provide a new outlook on the long-lasting storage of fear memory, suggesting that after the initial consolidation of memory, a secondary, systems consolidation-like phase of slow synaptic growth occurs, ultimately resulting in increased connectivity to the BLA fear circuit and an enhanced role of mPFC in fear memory retrieval.

## Materials and Methods

### Animals

All animal procedures were conducted in accordance with institutional guidelines and protocols approved by the Institutional Animal Care and Use Committee at The Scripps Research Institute and the University of California San Diego. FDC-Ribo mice were 5-7 months of age at the time of sacrifice. FDC mice for synaptophysin-venus experiments were 7-9 months of age at time of sacrifice. All mice were housed on a 12 hr light-dark schedule (light beginning at 6:00AM).

### FDC mouse production

The cfos-ddCre targeting construct was prepared by conventional cloning methods. Homology arms were PCR amplified from 129 genomic DNA (Phusion Hot Start II DNA Polymerase, Thermo-Scientific, cat. # F-549L) and cloned to flank a DD-Cre with 3′ SV-40 polyA cDNA and a downstream PGK-neo cassette. The 5′ homology arm consisted of 1823bp directly upstream of the cfos ATG. The 3′ homology arm consisted of 4259bp (85478035-85482293, Genbank accession number NC_000078.6), located 1576bp downstream of the cfos stop signal. After verification by sequencing, the linearized construct was electroporated into E14Tg2A ES cells, and clones with homologous recombination were recovered by long range PCR (Phusion Hot Start II DNA Polymerase, Thermo-Scientific, cat. # F-549L) (5′ arm: 5′-GTCTAACCCGGCTTGTCCTC-3′ and 5′-GTGTTCCTCTTGAACCAAGCCAGAT-3′; 3′ arm: 5′-tacccggtagaatgaagttcctatact-3′ and 5′-accactgctgtaactaagtcttcaaac-3′). Targeted ES cells were confirmed by radioactive southern blot and subsequently microinjected into BL/6 blastocysts. Germline transmission was detected by long range PCR with the same primers utilized for ES cell screening. cfos-ddCre heterozygous mice were fertile and did not have any obvious behavioral abnormalities. cfos-ddCre mice were crossed to a germline-active flpe expressing transgenic line to remove the PGK-neo cassette, and cassette removal was verified by PCR and sequencing (5′-GGACTTTCCTTCAGAATTGCTA-3′ and 5′-GGTCATGTTCTTCTTTGAGATGATAA-3′). All non-viral tdTomato visualization experiments utilized mice heterozygous for the cfos-ddCre allele and the ROSA26 Ai9 allele. Background recombination was indistinguishable between mice that harbored the pgk-neo cassette and mice that had it excised (data not shown), and therefore we used the initially obtained pgk-neo containing cfos-ddCre allele for all experiments in this manuscript.

### FDC mouse characterization procedures

For whisker barrel stimulation experiments, 4 week old male and female mice were anesthetized with isoflurane and whiskers were plucked under a dissecting microscope. Whiskers were removed on one side of the snout, and the other side kept intact for comparison. After whisker removal, mice were transferred to a new clean cage and allowed to recover for about 24 hours. Subsequently, mice were transferred to an “enriched” environment cage with new nesting materials and 2-3 L-shaped tubes, injected with 5mg TMP 2 hours later (equivalent to about 350ug/g TMP), and then sacrificed 14 days after TMP injection.

For all seizure induction experiments, group housed 7-9 week old male and female mice were weighed and separated into a single cage with the food and water removed. Mice were then injected intraperitoneally with kainic acid (Tocris biosciences, cat. # 0222) dissolved in PBS pH

7.4 to 25mg/kg. In most experiments, some mice did not survive after seizure and some mice did not have obvious seizures. Mice were only used for further processing if a seizure was observed. Seizures usually occurred about 5-10 min after kainic acid injection, and were characterized by severe tremor, especially of the forepaws, hunched back, and stiff tail. For seizure dose-response experiments, TMP was injected 1 hour after kainic acid injection. For time course of ligand induced labeling experiments, animals were injected with either TMP or TMP-lactate at various time points surrounding the injection of kainic acid. Animals were left undisturbed in between kainic acid and TMP injections. Animals were singly housed and sacrificed 7 days later for tdTomato visualization. For the TMP dose-response curve, vehicle mice were given 50ul DMSO in 500ul total PBS (10% DMSO). For the TMP time course, vehicle mice were given 20ul DMSO in 500ul PBS (4% DMSO).

### FDC-Ribo mouse production

To generate FDC-Ribo mice, RiboTag mice were obtained from The Jackson Laboratory (Stock #011029) and crossed to heterozygous FDC mice. FDC-Ribo mice used for RNAseq experiments were heterozygote FDC and homozygote for RiboTag. All other experiments were completed with mice heterozygous for both the RiboTag and FDC alleles.

### TMP injections

For FDC mouse characterization (Fig 1, Supplementary Fig 1), animals were administered TMP via intraperitoneal injection under the conditions described above. For all other experiments, mice were injected with TMP-lactate at 150μg/g 15 minutes following behavioral procedures.

### Behavioral experiments

Context fear conditioning was performed in Med Associates fear conditioning chambers with a black and white checkered background and peppermint scent (Dr. Bronner’s). Chambers were washed with water and 70% EtOH between sessions. Training consisted of 8 minutes total beginning with a 3 minute acclimation period followed by four 1s 1mA shocks with varying inter trial intervals from 45 – 85 seconds. Context exposure only was performed in the same context for 8 minutes total. Immediate shock training was a total of 8 minutes with four 1s 1mA shocks delivered within the first 14 seconds of the session (ITI 3s). All memory retrieval sessions were 5 minutes. Freezing behavior was measured using the Med Associates software in which immobility for 1s was considered a bout of freezing.

### Tissue dissection and immunoprecipitation

Mice were anesthetized with isoflurane and transcardially perfused with cold PBS. Brains were removed and placed in a stainless steel 1mm brain matrix (Stoelting, catalog #51386) and 2 brain slices were obtained spanning from bregma coordinates A/P +2.20 to +0.20. The mPFC (anterior cingulate, prelimbic, and infralimbic cortices) was removed using a 1mm tissue biopsy punch and immediately submerged in 800μl cold IP buffer [50mM Tris pH7.4; 100mM KCl; 12mM MgCl2; 1% NP-40; 100 μg/ml cycloheximide (Sigma); 1.0 mM DTT; 1x protease inhibitor (Promega); 1 mg/ml sodium heparin (Sigma); 0.2 units/μl RNasin (Promega)] in a dounce homogenizer. After homogenation, samples were centrifuged (10,000 × g for 15 min at 4°C), and 50μl supernatant was saved as “input” while remaining supernatant (lysate) was incubated with 7μl Mouse monoclonal anti-HA.11 (Ascites Covance MMS-101R) and 20μl Rabbit polyclonal anti-HA (Invitrogen cat#71-5500) at 4°C while rotating. After 4 hours, lysate was incubated with Pierce protein A/G magnetic beads (Invitrogen; washed in IP buffer first) overnight at 4°C while rotating. Tubes were then placed on magnetic stand and beads were washed 3 times with a high salt buffer [50mM Tris, pH 7.4; 300mM KCl; 12mM MgCl_2_; 1% NP-40; 100 μg/ml cycloheximide (Sigma); 1.0 mM DTT] before moving to the RNA isolation step.

### RNA isolation and cDNA synthesis

RNA was purified using Qiagen RNeasy Plus Micro kit (cat#74034) and checked for quality using an Agilent Bioanalyzer. Reverse transcription and cDNA synthesis/amplification was done using NuGen Ovation RNAseq V2 kit. qPCR was performed with QuantStudio5 (Thermofisher) with Taqman primer/probe sets and all values were normalized to Beta-2-Microglobulin (B2M).

### RNA sequencing

The 75bp reads are generated by the NextSeq Analyzer located at the Scripps DNA Sequencing Facility. The Genome Analyzer Pipeline Software (bcl2fastq/2.16.0.10) is used to perform the early data analysis of a sequencing run, which does the image analysis, base calling, and demultiplexing. The program called cutadapt is used to trim the adapter and low called basepair scores. For mRNA-Seq STAR/ 2.3.0 was used to align to the mouse genome reference (mm10). Differential expression analysis was performed using the R bioconductor package EdgeR. Pathway enrichment analysis was performed using the DAVID bioinformatics database^45^.

### Immunofluorescence

After completion of all experimental procedures, animals were anesthetized with isoflurane and transcardially perfused with PBS followed by 4% paraformaldehyde. Brains were postfixed in PFA overnight and sectioned by vibratome (Leica) at 70 μm. Free-floating brain sections were blocked in 10% normal goat serum/0.2% Triton × for two hours and subsequently probed with primary antibodies (see Supplementary Table 2 for antibody incubation details and dilutions) followed by three 10 minute washes in PBS. Sections were switched to Alexa Fluor secondary antibodies for 2 hours at room temperature followed by a 30 min incubation with DAPI (1:1000 in PBS). Sections were washed three times in PBS for 20 minutes, mounted, coverslipped and stored at 4° until imaged. For GAD67 staining, sections were permeabilized in PBS Triton × 0.1% for 30 minutes at room temperature prior to primary antibody incubation. For GAD67 and NeuN, sections had been stored in cryoprotectant at −20°C prior and were washed 3×5 minutes in PBS before staining.

### Microscopy and image analysis

Brain sections were imaged using an A1 Nikon Confocal microscope. All laser settings were held constant, and images were acquired blind to the experimental group. Cell counts and overlap quantification were completed using an ImageJ macro (available upon request) which identifies only cells that overlap with DAPI. For the reactivation experiment (Figure 2), cells co-expressing HA and cFos were considered reactivated (OL) and were normalized to the total cell population (HA+ cells + cFos+ cells – OL) to account for differences in cFos expression between groups. Reactivation is presented as a percent of control, in which control equals 100 percent. For synaptophysin-venus viral experiments, lower magnification (20x) analysis was completed as follows: ROIs were drawn around the BLA and striatum in ImageJ (three images per mouse), and the total area (μm^2^) was measured. Next, all images were thresholded to a constant (120, 255) to include only the brightest signal (by eye), and the thresholded area (μm^2^) was measured. Average μm^2^ signal/total μm^2^ was calculated for each mouse and normalized to its respective viral infectivity before averaging each group.

Higher magnification (40x) axon bouton analysis was completed for BLA and striatum (3 images per mouse) using the ImageJ analyze particles feature in which all images were thresholded to a constant (140, 255) and parameters for particle count were size=0-200; circularity=0.5-1.00. All analysis was completed blind to the experimental group.

### Surgeries and viral injections

FDC mice were anesthetized under an isoflurane (1-2%)/oxygen mix while head fixed in a stereotaxic apparatus (Kopf) before being bilaterally injected with a 1:3 ratio of DIO-Timer: DIO-Synaptophysin-venus (see below for AAV vector information) in mPFC with coordinates and volumes as follows: 300nl at +1.9 A/P; ±0.4 M/L; −2.1 D/V and 150nl at +1.0 A/P; ±0.5 M/L; −1.2 D/V. Injections were delivered at a rate of 100nl/min using a motorized pump, and the injection needle remained in place for 5 minutes both before and after injection. Veterinarian petrolatum ointment (Puralube) was used to prevent eyes from drying during surgery. When surgery was completed, the skin was closed with vicryl sutures (Ethicon VCO319) and 2.5 μg/g flunixin (Sigma-Aldrich, # 33586) was injected subcutaneously to reduce post-surgical pain. Mice were given at least 1 week to recover before behavioral experiments.

The DIO-timer (pAAV-CK2min-DIO-H2B-Timer; 4.5 × 10^13^ Genome Copies/ml) was produced in house, and the DIO-Syn-Venus (pAAV-Ef1α-DIO-Synaptophysin-Venus, 1.1 × 10^13^ Genome Copies/ml) was kindly provided by Anton Maximov at The Scripps Research Institute.

### Statistics

RNAseq differential expression analysis was done in R and all other statistics were performed in Prism Graphpad. Two-tailed unpaired student’s t-tests were used to compare experiments with two groups. Multiple group comparisions were analyzed using one- or two-way analysis of variance (ANOVA) with appropriate post hoc tests (noted in figure legends). All data presented as mean ± standard error of the mean (SEM).

## Author contributions

B.C.D and M.M. devised the main conceptual ideas behind this work. P.C. and L.S. designed the experiments included in Figure 1 and Supplemental figure 1. B.C.D. and M.M. designed all other experiments. P.C. completed all characterization experiments for the FDC mouse and corresponding analyses (Figure 1 and Supplemental figure 1). B.C.D. and S.P. characterized the FDC-Ribo mouse and optimized immunoprecipitation and RNA isolation procedures. B.C.D. performed behavioral experiments, molecular biology work, surgeries, immunostaining, microscopy and image analysis. L.M.C. assisted with experimental design, RNA isolation and contributed critically to scientific discussion, data interpretation and analysis. E.J.Y. assisted with immunostaining procedures. B.C.D and M.M. wrote the manuscript. P.C. and L.S. contributed to main text corresponding to Figure 1.

## Acknowledgements

We thank Ian R. Winchester and Yena J. Lee for assistance with the mouse colony and genotyping, and Natasha Weaver for administrative support during the years at The Scripps Research Institute. We thank Jason Keller for his contribution to the FDC mouse characterization.

## Supplemental information

**Supplemental Table 1.** FDC background recombination in 24 brain regions. Percentage of cells labeled/total cells at 4 and 8 weeks post birth in the absence of TMP, N=3-4/brain region, mean ± SEM.

| Brain Region | 4 weeks (% total) | 8 weeks (% total) |
| --- | --- | --- |
| **cerebellum** |  |  |
| molecular layer | 1.11 ± 0.56 | 1.49 ± 0.83 |
| granule cell layer | 2.15 ± 0.58 | 7.25 ± 1.48 |
| purkinje cell layer | 2.73 ± 0.30 | 7.37 ± 2.32 |
| **cortex** |  |  |
| primary somatosensory | 1.68 ± 0.34 | 4.10 ± 0.41 |
| primary motor | 0.35 ± 0.04 | 1.48 ± 0.26 |
| **hippocampus** |  |  |
| CA1 | 0.20 ± 0.07 | 1.03 ± 0.21 |
| dentate gyrus | 11.05 ± 1.88 | 43.82 ± 7.46 |
| **hypothalamus and amygdala** |  |  |
| anterior hypothalamus | 1.16 ± 0.41 | 2.45 ± 0.29 |
| arcuate nucleus | 3.56 ± 0.13 | 6.34 ± 0.73 |
| basolateral amygdala | 1.00 ± 0.33 | 2.89 ± 0.43 |
| BNST (posterior) | 1.03 ± 0.24 | 2.97 ± 0.31 |
| dorsal medial hypothalamus | 4.34 ± 1.26 | 5.52 ± 0.56 |
| medial amygdala (anterior) | 2.50 ± 0.40 | 5.91 ± 0.52 |
| medial amygdala (posterior) | 2.91 ± 0.53 | 5.90 ± 0.89 |
| medial preoptic nucleus | 1.38 ± 0.56 | 3.00 ± 0.50 |
| periaqueductal gray | 1.58 ± 0.27 | 2.43 ± 0.33 |
| posteromedial cortical amygdala | 4.06 ± 0.45 | 7.96 ± 0.36 |
| ventromedial hypothalamus | 3.59 ± 0.68 | 6.74 ± 0.54 |
| **basal ganglia and thalamus** |  |  |
| caudate putamen | 0.41 ± 0.03 | 0.71 ± 0.09 |
| lateral globus pallidus | 0.12 ± 0.9 | 1.59 ± 0.3 |
| ventral posteromedial nucleus | 2.93 ± 0.23 | 7.88 ± 2.35 |
| **olfactory bulb** |  |  |
| AOB mitral cell layer | 5.52 ± 0.39 | 6.56 ± 1.17 |
| MOB mitral cell layer | 9.11 ± 0.48 | 14.03 ± 0.92 |
| MOB granule cell layer | 10.28 ± 1.36 | 18.20 ± 1.29 |

**Supplemental Table 2.** Antibody information. Antibody catalog numbers, dilutions and incubation times for immunostaining.

| <u>Primary Antibody</u> | <u>Host</u> | <u>Type</u> | <u>Vendor</u> | <u>Catalog Number</u> | <u>Dilution</u> | <u>Antibody Incubation</u> |
| --- | --- | --- | --- | --- | --- | --- |
| Anti-GAD67 | Mouse | Monoclonal | EMD Millipore | MAB5406 | 1:500 | 48hr RT |
| Anti-cFos | Rabbit | Monoclonal | Cell Signalling | 2250 | 1:700 | O/N 4° |
| Anti-NeuN | Rabbit | Monoclonal | abcam | ab177487 | 1:1000 | O/N 4° |
| Anti-HA. 11 | Mouse | Monoclonal | BioLegend | 901513 | 1:1000 | O/N 4° |
| Anti-HA | Rabbit | Polyclonal | ThermoFisher | 71-5500 | 1:200 | O/N 4° |
| <u>Secondary Antibody</u> | <u>Host</u> | <u>Target</u> | <u>Vendor</u> | <u>Catalog Number</u> | <u>Dilution</u> | <u>Primary Used</u> |
| Alexa Fluor 488 | Goat | Mouse | abcam | ab150113 | 1:750 | Anti-GAD67; Anti-HA. 11 |
| Alexa Fluor 647 | Goat | Rabbit | abcam | ab150079 | 1:750 | Anti-cFos; Anti-Neun; Anti-HA |

**Supplemental Figure 1.**
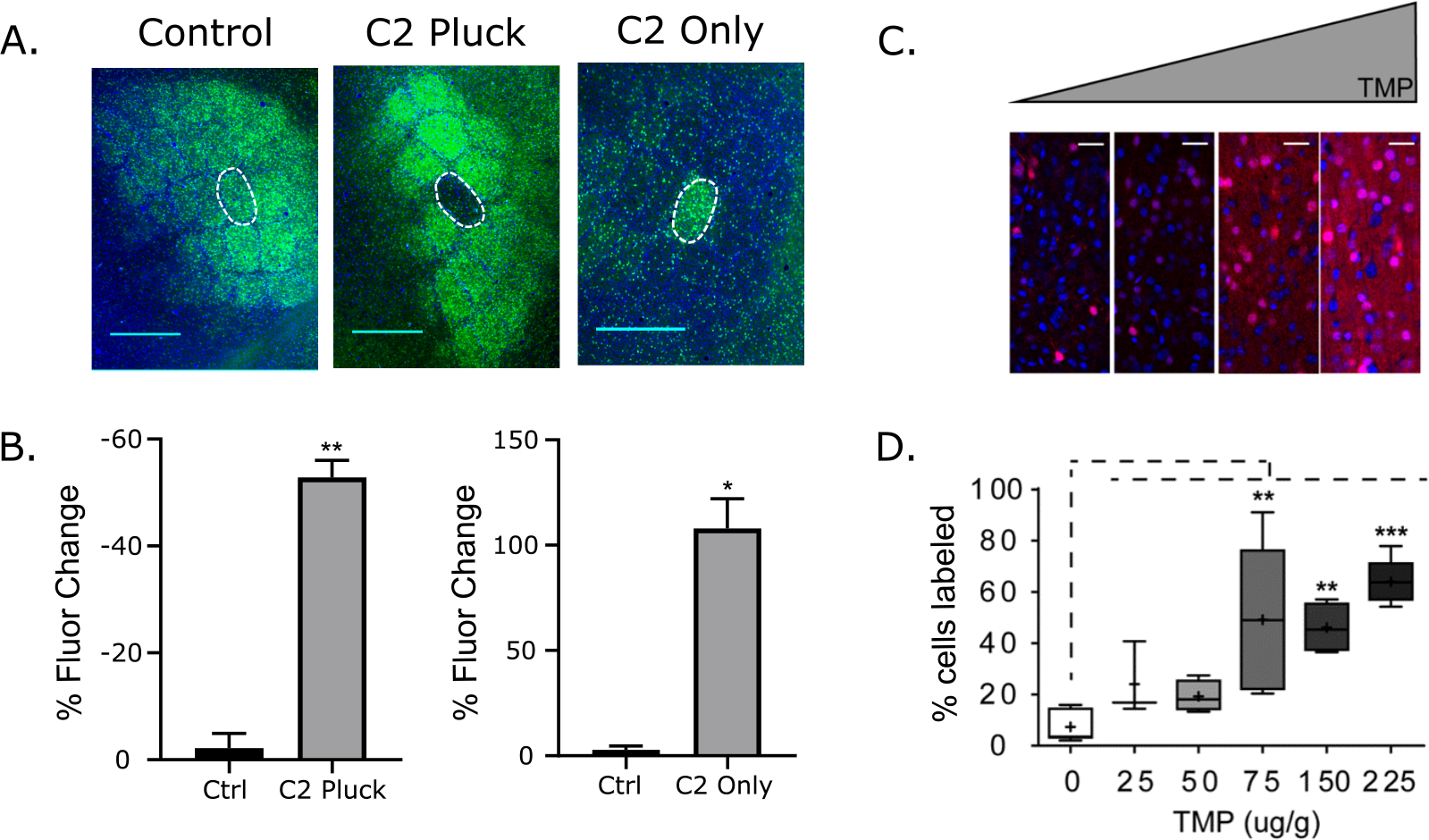
FDC mouse characterization. A) Representative images of whisker barrel experiment. No pluck control (left), C2 pluck (middle) and C2 pluck only (right). Scale bar =1mm B) Significant changes in fluorescence intensity of C2 pluck or C2 only groups relative to surrounding intact barrels. N=3; Mean ± SEM; students t-test, **p<0.01, *p<0.05. C) Varying concentrations of TMP were injected into mice 1 hour after kainic acid administration. Representative images of primary motor area, layer 1 to layer 2/3, after (left to right) 0μg/g, 25μg/g, 75μg/g, and 225μg/g TMP (Red=tdTomato; Blue=DAPI). D) Quantification shows a dose sensitive increase in labeling in primary motor area (N=3-7, box and whisker plot showing min and max as endpoints, mean represented as “+”, midpoint lines in box show median, samples compared to 0μg/g, Dunnett’s multiple comparisons test, *p<0.05, **p<0.01, ***, p<0.005).

**Supplemental Figure 2.**
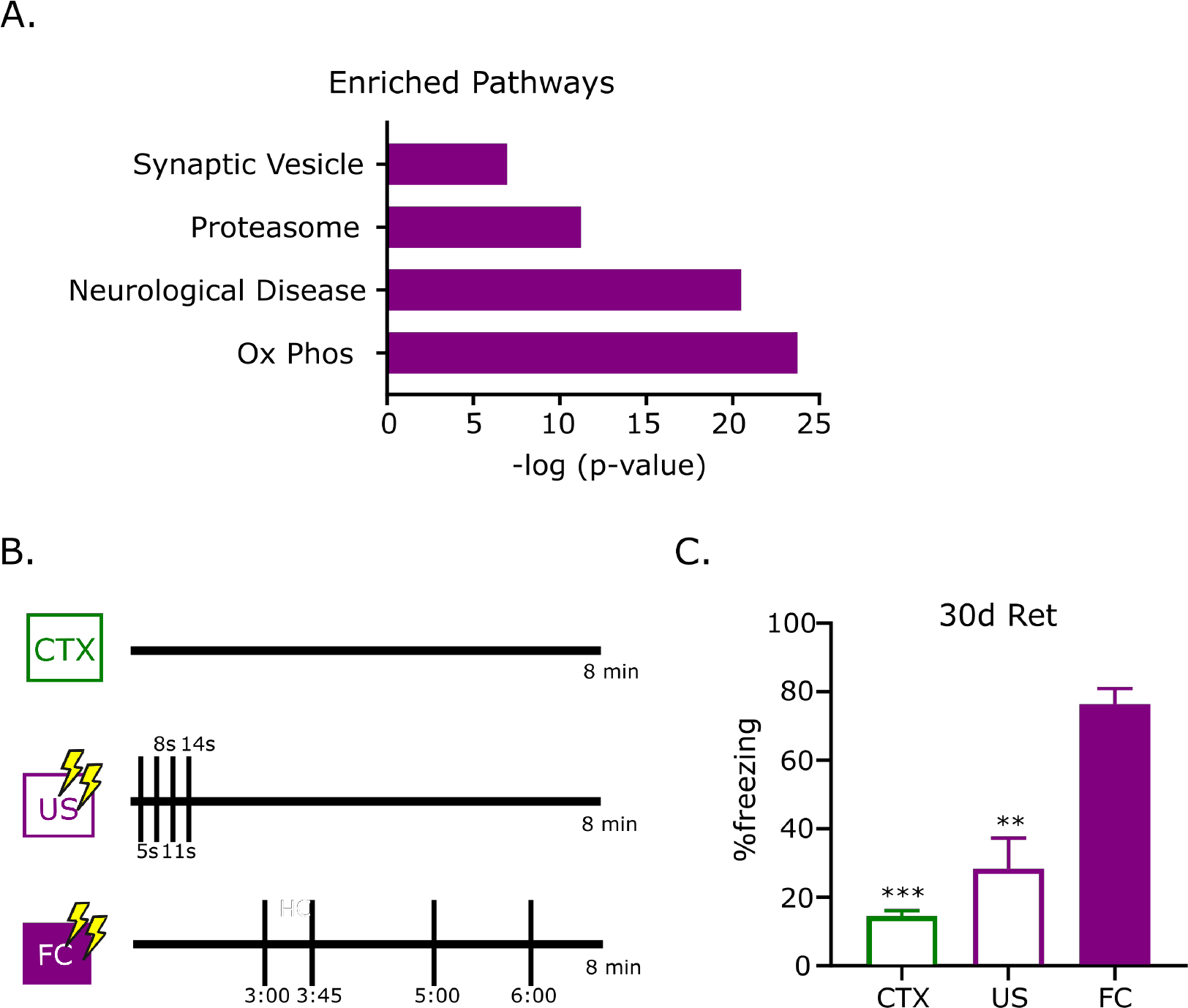
Pathway enrichment and behavioral data for gene expression experiments. A) Significantly enriched pathways in FC mice (Ox Phos refers to oxidative phosphorylation). Enrichment analysis completed using the DAVID open source bioinformatics tool. B) Experimental design. Mice were exposed to the same context for 8 minutes. The context only group (CTX) did not receive any electrical shocks. The immediate shock group (US) was delivered four 1mA 1 second shocks (denoted by vertical bars) at the beginning of the session with an inter trial interval (ITI) of 3 seconds. The fear conditioned group (FC) received four shocks 1mA after a 3 minute acclimation period with an average ITI of 60 seconds. C) Mice were returned to the training context 30 days later without any electrical shocks. Both CTX and US groups froze significantly less than FC group but were not significantly different from each other. N=4. Mean ± SEM. Tukey’s multiple comparisons test **p<0.01 ***p<0.001. All asterisks represent significant differences relative to FC.

**Supplemental Figure 3.**
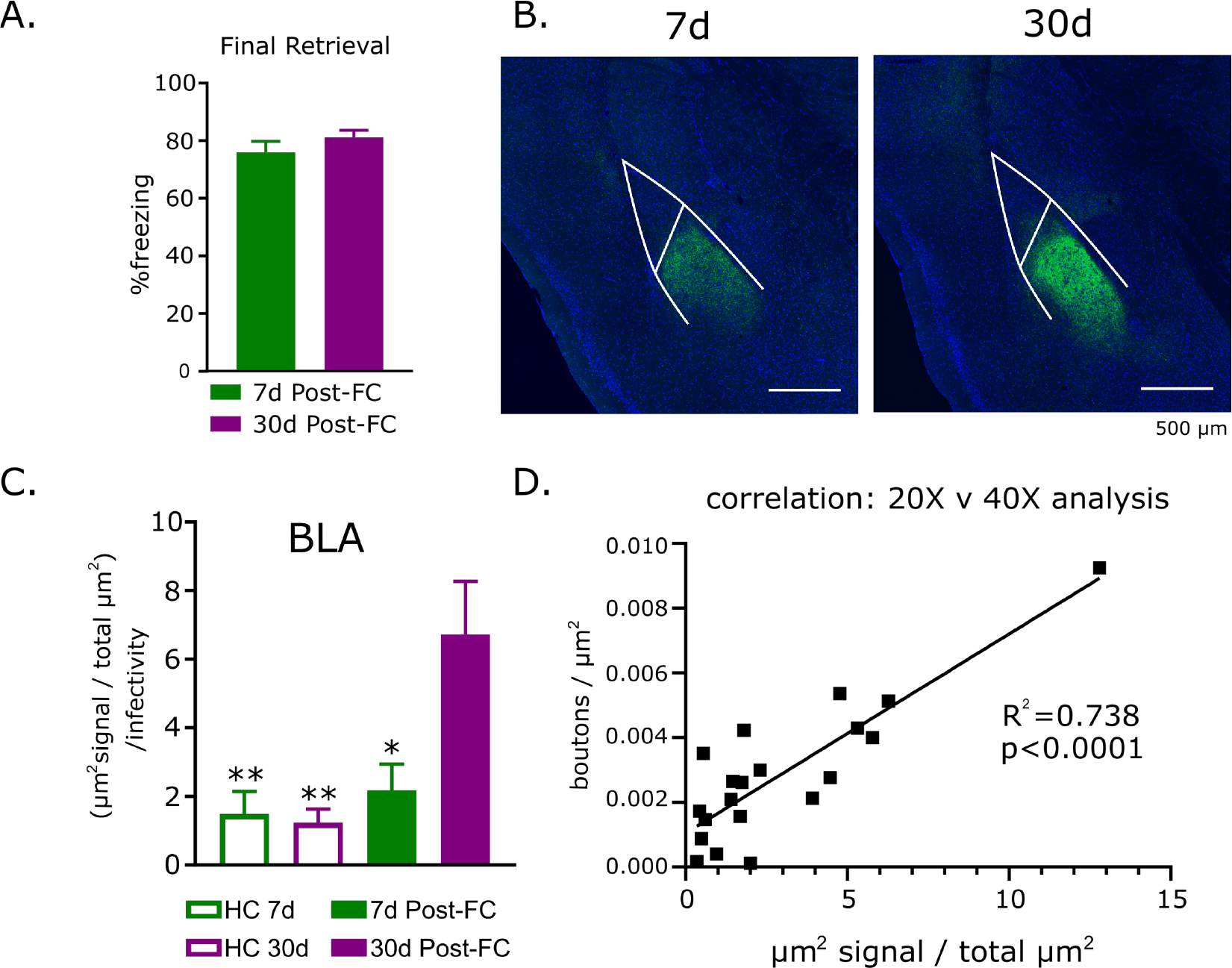
Behavioral data and lower magnification analysis of learning-activated mPFC-BLA projection neurons. A) No significant differences in freezing during the final retrieval trial between 7d Post-FC and 30d Post-FC. N=5-6; Mean ± SEM. B) Representative images of BLA at 20x magnification in 7d post-FC mice (left) and 30d post-FC mice (right). C) Significantly more synaptophysin-venus expression [(μm^2^ signal/total BLA μm^2^)/infected cells] in 30d post-FC group relative to 7d post-FC (*p=0.0156) as well as 7 d (**p=0.0076) and 30d (**p=0.0084) HC controls. N=5-6. Mean ± SEM. Tukey’s multiple comparisons test; all p-values are relative to 30d post-FC group. D) 20X [(μm^2^ signal/total BLA μm^2^)/infected cells] analysis is significantly correlated with 40X [boutons/μm^2^/infected cells]. See methods for detailed analysis procedures.

